# Metabolic Reaction Network-based Recursive Metabolite Identification for Untargeted Metabolomics

**DOI:** 10.1101/305201

**Authors:** Xiaotao Shen, Xin Xiong, Ruohong Wang, Yandong Yin, Yuping Cai, Zaijun Ma, Nan Liu, Zheng-Jiang Zhu

**Affiliations:** Interdisciplinary Research Center on Biology and Chemistry, Shanghai Institute of Organic Chemistry, Chinese Academy of Sciences, Shanghai, 200032 P. R. China; University of Chinese Academy of Sciences, Beijing, 100049 P. R. China

**Author notes:** **Corresponding Author** Correspondence should be addressed to Z.J.Z.

## Abstract

Metabolite identification is a long-standing challenge in untargeted metabolomics and a major hurdle for functional metabolomics studies. Here, we developed a metabolic reaction network-based recursive algorithm and webserver called MetDNA for the large-scale and unambiguous identification of metabolites (available at http://metdna.zhulab.cn). We showcased the versatility of our workflow using different instrument platforms, data acquisition methods, and biological sample types and demonstrated that over 2,000 metabolites could be identified from one experiment.

---

A major goal of LC-MS-based untargeted metabolomics is to provide mechanistic insights by systematically characterizing metabolic changes in relevance to the physiological and pathological status at the system level^1–5^. Yet large-scale, unambiguous identification of metabolites remains a challenging task in the field of untargeted metabolomics^3–5^. One widely used strategy in the identification of metabolites is to match experimental tandem mass spectrometry data (MS2 spectrum) with those from the standard spectral library^6^. This strategy, however, has a substantial limitation given the facts that only less than 10% of known metabolites in HMDB^7^ and METLIN^8^ have standard MS2 spectra, and that expanding the spectral library is limited by the availability of chemical standards for many metabolites. Moreover, this strategy also suffers from having a poor standardization protocol for curating spectral libraries, due to uncharacterized spectral variations across different instruments and laboratories^9^. Despite that many efforts have been made to predict the MS2 spectra *in silico*^10–14^, accuracy still requires substantial improvement^15^.

Here, we developed **Met**abolite identification and **D**ysregulated **N**etwork **A**nalysis (MetDNA) software, which implemented a metabolic reaction network (MRN)-based recursive algorithm for metabolite identification (Fig.1). This algorithm allows the large-scale metabolite identifications with a small tandem spectral library. In cellular metabolism, different metabolites can be transformed into metabolite products by appropriate enzymatic reactions. We define a reaction pair (or reactant pair, RP) by pairing a substrate metabolite and its product metabolite with similar chemical structures^16^. We thus retrieved all the metabolite RPs from the KEGG database (9,603 RPs and 7,639 metabolites, http://www.kegg.jp/) and constructed a metabolic reaction network (Fig. 1a). In the MRN, one node represents one metabolite, and two metabolites connected by an edge represent the reaction pair or so-called neighbor metabolites. In tandem MS, the fragmentation pattern of a metabolite depends on its chemical structure. Therefore, due to their structural similarities, neighbor metabolites tend to share similar MS2 spectra (Fig. 1a and **Supplementary Fig. 1**). Using this principle, MetDNA was deployed to identify metabolites without existing MS2 spectra in the spectral library. We tested this approach by first characterizing a few metabolites using a small tandem spectral library, and utilized their experimental MS2 spectra as surrogate spectra for identifying their neighbor metabolites (Fig. 1b). This approach facilitates metabolite identifications without expanding the spectral library. Importantly, the reiterated application of this surrogate principle allows significant and progressive expansion of identified metabolites with the MRN through our recursive algorithm (Fig. 1c).

**Figure 1.**
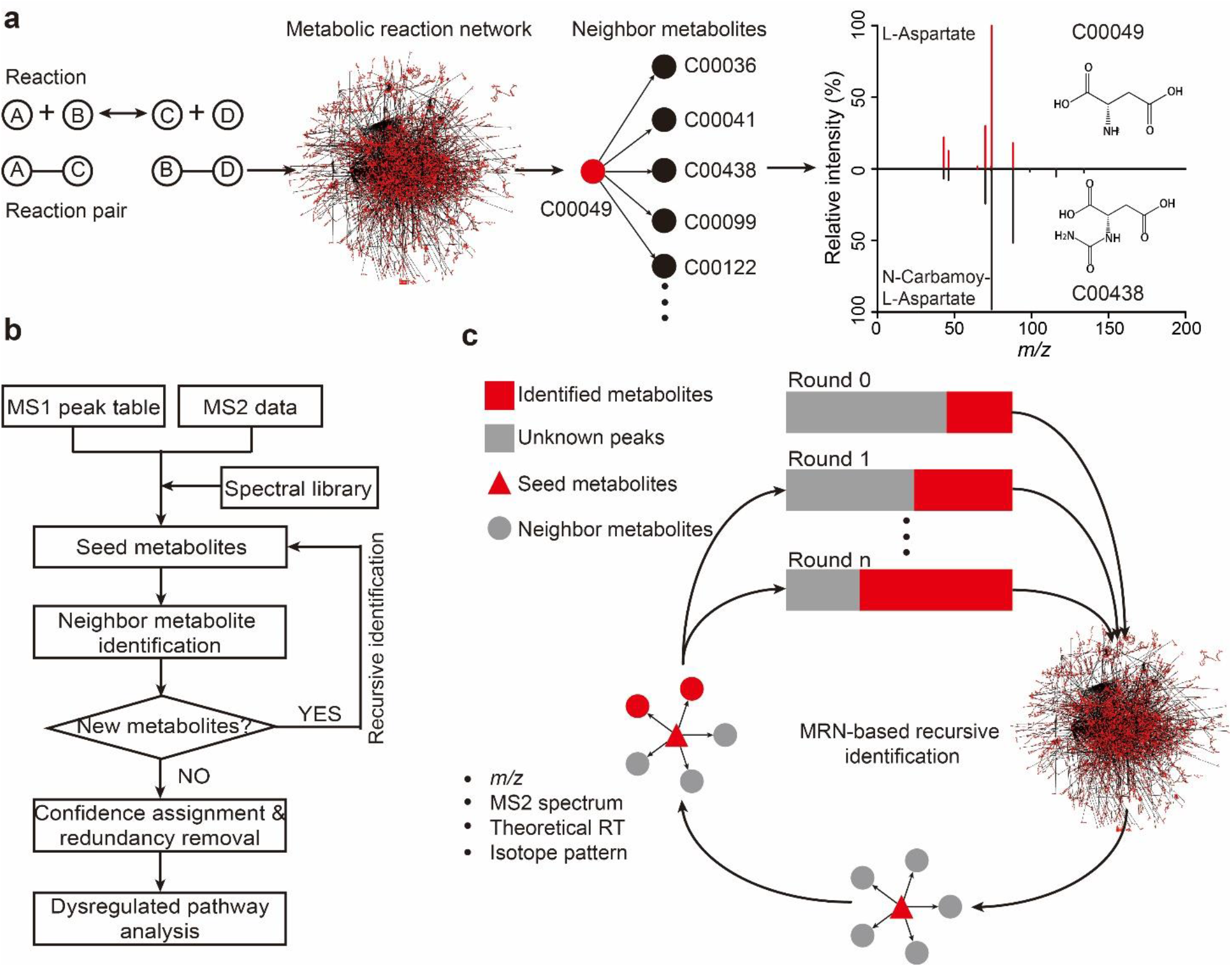
Metabolic reaction network (MRN) and the MetDNA workflow. (**a**) The construction of MRN using the reaction pairs retrieved from KEGG. An example is given to show that structurally similar neighbor metabolites have high similarity within MS2 spectra. (**b**) The overall workflow for MetDNA: (1) import of MS1 peak table and MS2 data; (2) identification of initial seed metabolites; (3) identification of neighbor metabolites; (4) MRN-based recursive identification; (5) confidence assignment and redundancy removal; and (6) dysregulated pathway analysis. (**c**) Illustration of MRN-based recursive identification. Seed metabolites (red triangle) from round 0 are first mapped to MRN and all neighbor metabolites are retrieved (gray circle). The neighbor metabolites are then identified by using the matching *m/z*, surrogate MS2 spectrum, theoretical RT and isotope pattern with unknown peaks (gray square, round 1). The recursive identification runs until there are no new identified metabolites (round n).

We next showcased the MetDNA workflow using untargeted metabolomics data acquired from aging samples of the fruit fly (*Drosophila melanogaster*; 3-day *vs*. 30-day; Fig. 2 and **Online Methods**). The fruit fly has been an important model organism for aging studies. However, the metabolic changes during adult lifespan of fruit flies remain to be explored. For the positive mode dataset, we detected a total of 18,320 MS1 peaks using XCMS^17^. Next we imported the generated MS1 peak table and MS2 data files into MetDNA. We first identified 115 metabolites using the standard spectral library. Among 115 metabolites, we selected 113 metabolites with KEGG IDs as the initial seed metabolites to map the MRN and retrieved their neighbor metabolites (Fig. 1c and Fig. 2a). At the first round, 581 neighbor metabolites were retrieved. Specifically, 145 of the 581 metabolites were identified by matching the calculated *m/z*, theoretical retention times (RTs) and surrogate MS2 spectra from the seed metabolites with the experimental data (Fig. 2a and **Supplementary Fig. 2**). We next selected 103 out of the 145 metabolites as the seed metabolites for the second round of identification. This recursive identification process was reiterated for 20 rounds, at which step no new metabolites could be identified. Finally, we evaluated the confidence of the metabolite identifications and removed redundant identifications (see **Online Methods**). This analysis identified a total of 1,496 metabolites from the positive mode (Fig. 2a and **Supplementary Figs. 2 and 3**). We next repeated this analysis for the negative mode and identified a total of 1,538 metabolites through 14 rounds of recursive identification (Fig. 2b and **Supplementary Figs. 4** and **5**).

**Figure 2.**
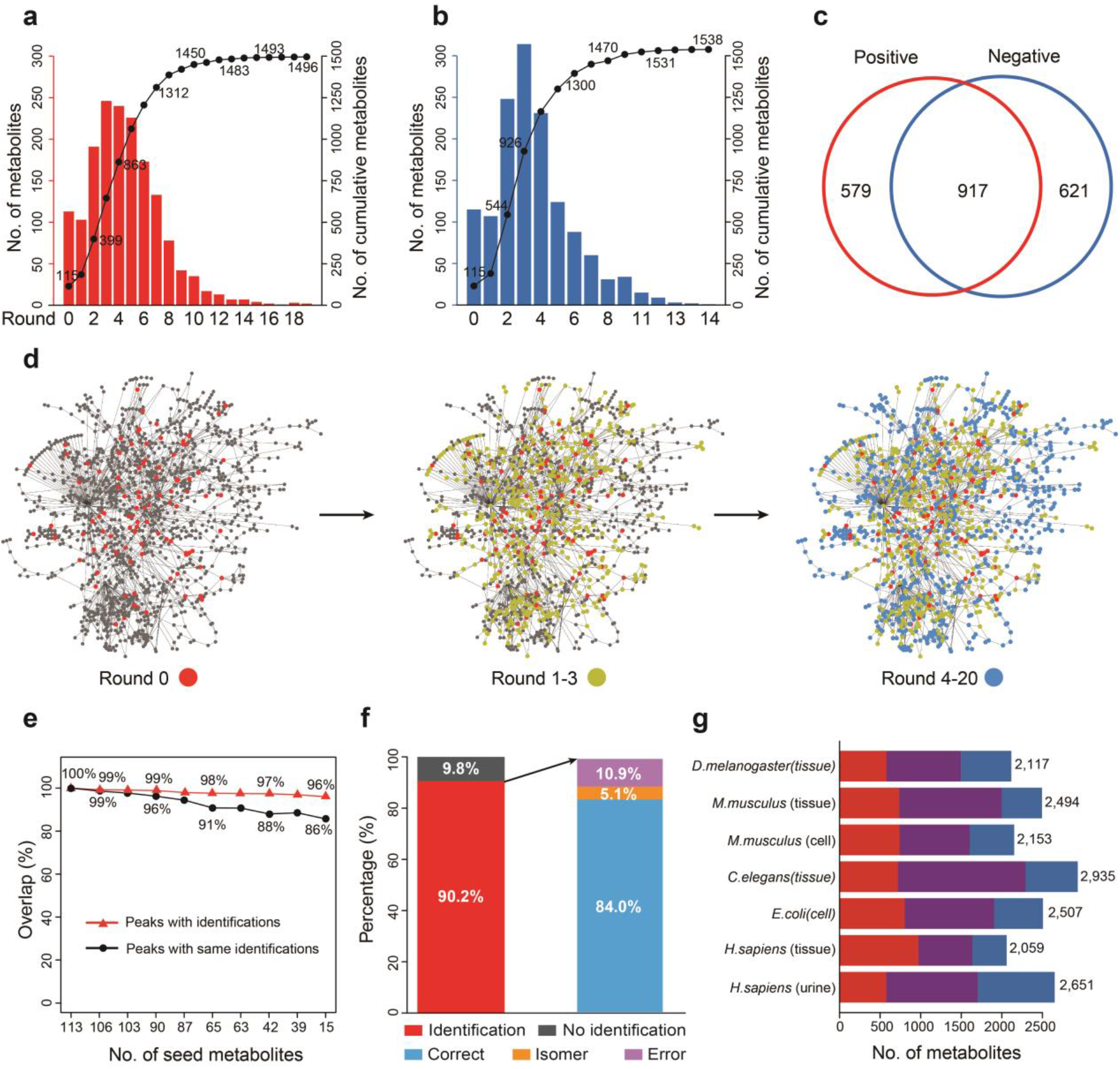
Metabolic reaction network (MRN)-based recursive metabolite identification. (**a**, **b**) The numbers of identified metabolites in aging fruit fly datasets: (**a**) positive and (**b**) negative modes. The left y-axis of the plot is the number of identified metabolites in each round, and the right y-axis of the plot is the cumulative number of identified metabolites. (**c**) A Venn diagram showing the overlap of identified metabolites in positive (red) and negative modes (blue). (**d**) Network diagrams demonstrating the distributions of identified metabolites in round 0 (left, red), rounds 1-3 (middle, yellow) and rounds 4-20 (right, blue). (**e**) The influence of the initial seed number on the identifications. Red triangles represent the overlap percentages of MS1 peaks with identifications using different initial seed metabolites. Black circles represent the overlap percentages of MS1 peaks with the exact same identifications using different initial seed metabolites. In both cases, the identification result using all 113 seed metabolites is used as a control. (**f**) Validation of metabolite identifications in the positive mode of aging fruit fly dataset. (**g**) The numbers of identified metabolites in different biological species and samples. The red color represents the results from the positive mode. The blue color represents the results from the negative mode. The purple color represents the overlap between the positive and negative modes.

To sum, a total of 2,117 metabolites were identified from a single experiment (Fig. 2c). Notably, majority of the metabolites were identified from the first eight recursive rounds, which accounted for more than 85% of total identified metabolites. Importantly, as the recursive identification progressed, we observed that the confidence of identified metabolites remains similar (**Supplementary Fig. 6**). Here we use an initial seed metabolite (adenosine diphosphate, ADP) as an example to demonstrate that MetDNA identified 4 metabolites in a recursive manner: adenosine diphosphate (AMP), inosine monophosphate (IMP), N6-(1, 2-Dicarboxyethyl)-AMP and inosine (**Supplementary Fig. 7**). The structures of these 4 metabolites were further validated using chemical standards to prove the accuracy of MetDNA (**Supplementary Fig. 7**). To conclude, our data indicates that the application of MetDNA provides a substantial expansion of identified metabolites from a just small number of initial seed metabolites (Fig. 2d).

We further evaluated how the number of initial seed metabolites impacts the identification result. Using the fruit fly dataset, we randomly selected a small fraction of seed metabolites as the initial seeds for recursive identification. Interestingly, the use of 15 metabolites as seeds was sufficient to identify similar number of metabolites compared to that of 113 seed metabolites (Fig. 2e and **Supplementary Fig. 8**). In addition, 86% of the identified peaks had the exactly same identifications as those derived from 113 seed metabolites (Fig. 2e). When using more initial seed metabolites, the overall confidence levels were slightly elevated and consequently the redundancy in the final result was slightly reduced (**Supplementary Fig. 8**). We observed a similar result from metabolomics dataset of aging mouse samples (**Supplementary Fig. 9**), suggesting that the overall performance of MetDNA analysis has been little influenced by the number of initial seeds. Therefore, the use of a small tandem spectral library to identify initial seed metabolites is sufficient.

How reaction steps involved determine the identification result was another critical aspect to be evaluated. Indeed, the inclusion of two or more reaction steps when retrieving the neighbor metabolites significantly increased the number of identified metabolites compared to one reaction step, while maintaining similar levels in confidence and redundancy (**Supplementary Fig. 10**). Similar results were obtained using aging datasets from fly and mouse (**Supplementary Fig. 10**).

To validate the identifications obtained from MetDNA, we first confirmed the chemical structures of the initial seed metabolites in the fruit fly dataset using commercial chemical standards (**Supplementary Tables 1** and **2**), which are considered as the Level 1 identifications according to MSI^18^. Using the initial seed metabolites, we designed a validation strategy (**Supplementary Fig. 11** and **Online Methods**). Specifically, 30% of the metabolites were randomly selected as the seeds, while the remaining 70% were used for the validation. The validation process was repeated 10 times. Finally, 90.2% of the metabolites in the validation metabolites were successfully identified (Fig. 2f). The overall percentage of the correct identification was 84.0%, whereas isomer and erroneous identifications were only 5.1% and 10.9%, respectively (Fig. 2f and **Supplementary Data 1**). Similar results were also obtained for negative mode data (**Supplementary Fig. 11** and **Supplementary Data 1**). We also repeated the validation process using additional independent datasets and found that the identifications from MetDNA were highly accurate (**Supplementary Fig. 11**, **Supplementary Tables 3** and **4**, and **Supplementary Data 2**).

We further tested MetDNA workflow with additional biological datasets. We found that over 2,000 metabolites could be identified from a variety of biological samples by this approach (Fig. 2g, and **Supplementary Tables 5** and **6**). These datasets covered five species and seven different sample types. The data acquisition used three different instrument platforms (Sciex TripleTOF, Agilent QTOF and Thermo Orbitrap) and the MS2 data were obtained using different acquisition methods such as data dependent acquisition (DDA), data independent acquisition (DIA) or targeted MS2 acquisition (**Supplementary Note 1**). Thus, MetDNA provides a platform-independent and versatile software tool.

In addition to metabolite identification, MetDNA can also perform dysregulated pathway analysis. In the aging fruit fly dataset, the statistical analysis discovered 875 dysregulated peaks with identifications (Student’s *t*-test, FDR-corrected *P*-values < 0.01, Fig. 3a and **Supplementary Data 3**). The pathway enrichment analysis identified 24 dysregulated metabolic pathways (Hypergeometric test, *P*-values < 0.05, Fig. 3b and **Supplementary Data 4**)^19^. Most of the dysregulated pathways that showed age-dependent regulation were associated with amino acid and sugar metabolism (Fig.3c and **Supplementary Fig. 12**).

**Figure 3.**
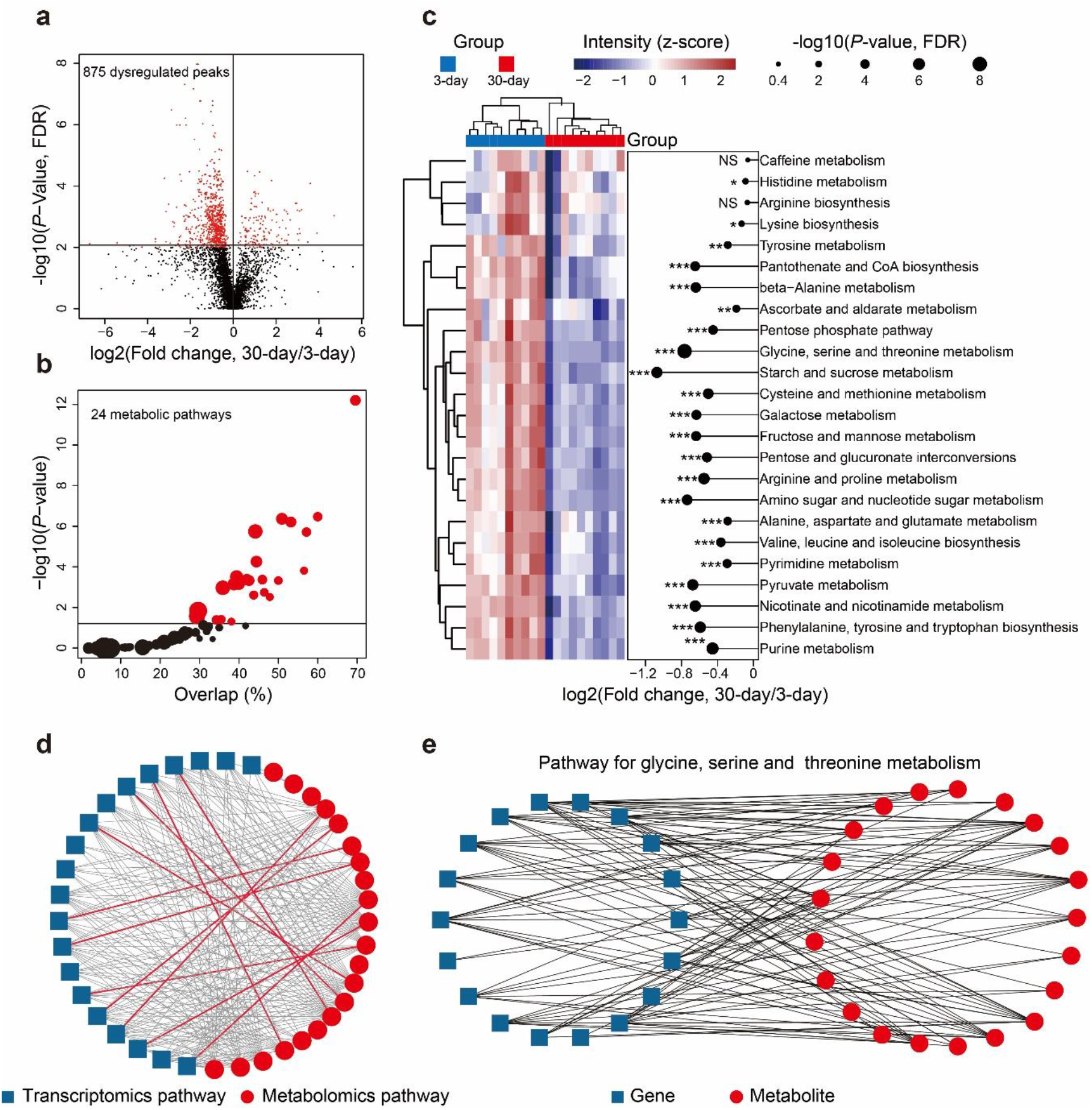
Dysregulated pathway analysis and multi-omics analysis. (**a**) A volcano plot showing the dysregulated metabolites (red dots) in the aging fruit fly dataset (Student’s *t*-test, FDR-corrected *P* values < 0.01). (**b**) The pathway enrichment analysis. The x-axis represents the overlap of identified metabolites for each pathway. The y-axis represents the −log10(*P*-value). The dot sizes represent the metabolite numbers in the pathway. The red color indicates the enriched pathways (Hypergeometric test, *P*-values < 0.05). (**c**) A heatmap showing the expression levels of 24 enriched metabolic pathways in aging fruit flies. The color indicates the z-score transformed intensity of the pathway. (**d**) The correlative network for pathways in metabolomics and transcriptomics. One node represents one pathway. The connections between the nodes were established by Pearson correlation (Student’s *t*-test, *P*-values < 0.05). The gray lines indicate the connections between two different pathways, and the red lines indicate the connections between the same pathways. (**e**) The correlation network of genes and metabolites in the pathway of glycine, serine and threonine metabolism. One node represents one gene or one metabolite. The black lines indicate the connections between genes and metabolites, and they were established by Pearson correlation (Student’s *t*-test, *P*-values < 0.01 and absolute Pearson correlation values > 0.7).

With substantially enlarged metabolomics data, we are now able to perform a multi-omics analysis by integrating the metabolomic data with transcriptomic data. A total of 74 and 51 metabolic pathways were quantitatively profiled using transcriptional and metabolomic analyses, respectively (**Supplementary Data 5, 6, 7** and **8**). A correlation network was constructed to demonstrate the high consistency in the changes between metabolites and gene expression^20^ (Fig. 3d). Eleven out of 24 overlapped metabolic pathways were highly correlated (red lines in Fig. 3d; Student’s-*t* test, *P*-values < 0.05). The metabolism of glycine, serine and threonine is a prominent example (KEGG ID: dme00260) of highly correlated at the metabolite and gene levels, as shown by a correlation network with 114 edges comprising 18 genes and 21 metabolites (Fig. 3e, **Supplementary Figs. 13** and **14**). The high correlation between gene expression and metabolites further validated the high confidence of metabolite identifications using MetDNA.

In conclusion, a new software tool named MetDNA has been developed for the large-scale and unambiguous identification of metabolites in untargeted metabolomics. MetDNA significantly increases the number of identified metabolites by using a MRN-based recursive strategy. In addition, MetDNA can also perform a dysregulated pathway analysis and provide quantitative evaluations for pathways and metabolites. By expanding the metabolite identification and integrating metabolomic and genomic data, MetDNA can become a powerful tool for integrative multi-omics analysis. MetDNA is available as a webserver with user-friendly interfaces (http://metdna.zhulab.cn/) and is free for non-commercial use.

## METHODS

Methods and any associated references are available in the online version of the paper.

*Note: Any Supplementary Information and Source Data files are available in the online version of the paper*.

## ACKNOWLEDGMETS

The work is financially supported by National Natural Science Foundation of China (Grant No. 21575151 to Z.J.Z) and the Thousand Youth Talents Program from Government of China (Z.J.Z). The work is also partially supported by Chinese Academy of Sciences Major Facility-based Open Research Program.

## AUTHOR CONTRIBUTIONS

Z.J.Z. and X.S. conceived the idea and designed the algorithm and software. X.S. developed the MetDNA program. X.X. developed the webserver. Y.Y. contributed to part of MetDNA code. R.W., Y.C., and X.S. performed the sample preparation, data acquisition and data processing. Z.M. and N.L. contributed to the RNA-seq data. X.S. and R.W. performed the data analysis. Z.J.Z., X.S., X.X. and R.W. tested and debugged the program and webserver. Z.J.Z. and X.S. wrote the manuscript. Z.J.Z. supervised the project.

## ONLINE METHODS

### Metabolic reaction network (MRN)

The metabolic reaction network is a network for metabolite-to-metabolite-based enzymatic reactions, and it was constructed using the KEGG database (http://www.kegg.jp/). In one metabolic reaction, the substrate and product metabolites are paired according to their structural similarity, and defined as one reaction pair (or reactant pair, RP)^16^. There are over 15,000 RPs defined in the KEGG. Here, we selected 9,603 RPs with specific metabolites and discarded the remaining generic reactions, symbolic reactions, or reaction pairs that only had small molecules such as oxygen and water. All the selected RPs were then combined to construct the MRN using the R package “*igraph*”. In the MRN, one node represents one metabolite, and one edge represents one reaction pair. Two metabolites connected by one edge indicate that they are neighbor metabolites. Finally, the MRN contains 7,639 metabolites (nodes) and 9,603 reaction pairs (edges) in total. Detailed information of all the reaction pairs in the MRN is provided in **Supplementary Data 9**.

### Data import

MetDNA requires the import of a MS1 peak table (.csv format) and MS2 data files (.mgf or.msp format). The MS1 peak table is a list of metabolic peaks with annotated *m/z* and retention times (RTs). The MS1 peak table is generated from the raw MS files using common peak picking software such as XCMS^17^ and MS-DIAL^21^. MS2 data files (.mgf format) are converted from MS raw files using ProteoWizard (version 3.0.6150, http://proteowizard.sourceforge.net/). MS2 data from different data acquisition methods such as data dependent acquisition (DDA), data independent acquisition (DIA) or targeted MS2 acquisition are all supported. If MS-DIAL is used for peak picking in DIA data, the generated MS2 data files (.msp format) are also supported by MetDNA. The details and examples about the generation of the MS1 peak table and MS2 data files are provided in **Supplementary Note 2**.

### Identification of initial seed metabolites

With the imported MS1 peak table and MS2 data, MetDNA first matches the MS1 peaks with MS2 spectra according to their *m/z* (± 25 ppm) and RT (± 10 s) values. If one MS1 peak matches multiple MS2 spectra, the most abundant MS2 spectrum is selected. In MetDNA, the intensities of the top 10 abundant fragment ions are summed up to represent the abundance of MS2 spectra. If one uses MS-DIAL for the peak picking and outputting MS2 data (.msp format), then the MS1-MS2 match step is skipped by MetDNA. The generated MS1/MS2 pairs are matched with our in-house standard spectral library for metabolite identification. The match tolerance for the MS1 *m/z* value is set as ± 25 ppm. The dot-product (DP) function^22^ is used to score the similarity between the experimental spectrum and the standard spectrum in the library (eq. 1). The DP score ranges from 0-1, from no match to a perfect match. The intensities of the fragment ions in the MS2 spectra are rescaled so that the highest fragment ion is set to 1.

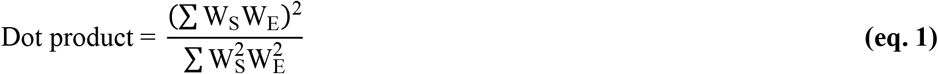

where weighted intensity vector, W = [relative intensity of fragment ion]^n^[*m/z* value]^m^, n = 1, m = 0; S = standard, and E = experiment. DP scores from both forward and reverse matches are generated. Metabolite identifications with DP scores in either forward or reverse matches larger than 0.8 are kept. Usually, 50-200 metabolites are identified using the standard spectral library. The identifications are further filtered using the theoretical RTs of the metabolites (± 30%). The generation of the theoretical RTs is provided below. The remaining identifications are used as “initial seed metabolites” in round 0 for MRN-based recursive annotation. The seed metabolites are defined as the identified metabolites that provide their MS2 spectra as surrogate MS2 spectra for the identification of their neighbor metabolites.

### Annotation of isotope and adduct peaks

First, the isotope peaks of seed metabolites are annotated by MetDNA. For each seed metabolite, the program calculates the theoretical *m/z* and the relative intensities of isotope peaks from the formula using the binomial and McLaurin expansion (R package “*Rdisop*”). Only four isotope peaks ([M] to [M+4]) are calculated. The generated isotope peaks of seed metabolites are used to match all the MS1 peaks in the MS1 peak table according to the *m/z*, RT and relative intensity. The default tolerances for the *m/z*, RT and relative intensity are set as ± 25 ppm, ± 3 s, and ± 500%, respectively. The large tolerance for the relative intensity match is due to the inaccuracy of the experimental intensity ratios for the isotope peaks^23^.

The matched isotope peaks are assigned the annotation scores (Score_iso_) shown below.

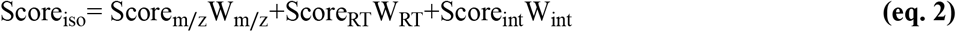

where Score_m/z_ represents the *m/z* match score and is calculated as follows:

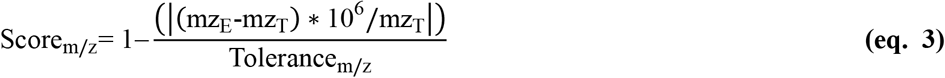

mz_E_ and mz_T_ are the experimental *m/z* and theoretical *m/z*, respectively. The Tolerance_m/z_ represents the tolerance for the *m/z* match with a default value of 25 ppm.

Score_RT_ represents the retention time match score and is calculated as follows:

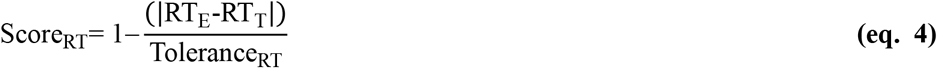

RT_E_ and RT_T_ are the experimental RT and theoretical RT, respectively. The Tolerance_RT_ represents the tolerance of RT match with a default value of 3 seconds.

Score_int_ represents the relative intensity match score and is calculated as follows:

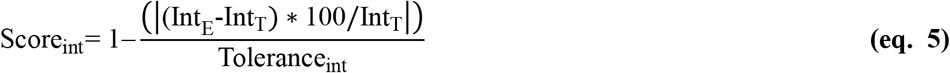

Int_E_ and Int_T_ are the experimental relative intensity and theoretical relative intensity, respectively. The Tolerance_int_ represents the relative intensity match tolerance with a default value of 500%.

W_m/z_, W_RT_ and W_int_ represent the weights of Score_m/z_, Score_RT_ and Score_int_, respectively, and the default values are 0.45, 0.45 and 0.1, respectively.

Second, MetDNA calculates the possible *m/z* values for the adduct peaks of seed metabolites. The adduct table for different LC separations (HILIC or RP) and polarities (positive and negative) are listed in **Supplementary Table 7**. All the possible adduct peaks of the seed metabolites are then matched to the MS1 peak table according to their *m/z* and RT. The default tolerances for the *m/z* and RT are ± 25 ppm and ± 3 s, respectively. The annotations of the adduct peaks are also assigned annotation scores (Score_adduct_) as follows.

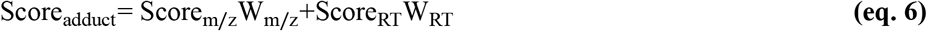

where Score_m/z_ represents the *m/z* match score and is calculated as indicated in eq. 3. Score_RT_ represents the retention time match score and is calculated using eq. 4. W_m/z_ and W_RT_ represent the weights of Score_m/z_ and Score_RT_, respectively, and the default values are 0.5, and 0.5, respectively.

Third, after the annotation of the adduct peak, the isotope peak annotation for each adduct peak is also performed using the same procedures described above.

### Identification of neighbor metabolites

Two metabolites in one reaction pair are neighbor metabolites. The seed metabolites provide their experimental MS2 spectra as surrogate spectra to identify their neighbor metabolites. First, the initial seed metabolites are mapped to MRN and retrieve their neighbor metabolites from MRN. The reaction step is defined as the number of reactions between two metabolites. All the neighbor metabolites with 1 reaction step are retrieved. The MS2 spectra of the seed metabolites are assigned to their corresponding neighbor metabolites as surrogate MS2 spectra. All possible adduct peaks of the neighbor metabolites are calculated according to the LC (HILIC or RP) and polarity. For each neighbor metabolite, the generated theoretical *m/z*, theoretical RT (from RT prediction) and surrogate MS2 spectrum are matched to the experimental MS1 *m/z* (default tolerance: ± 25 ppm), RT (default tolerance: ± 30%), and MS2 spectrum (default tolerance: 0.5). If one seed metabolite leads to no identification of a neighbor metabolite with 1 reaction step, then neighbor metabolites with 2 reaction steps are further retrieved. The default value of the maximum reaction step is 3. The identifications of neighbor metabolites are assigned the identification scores shown below.

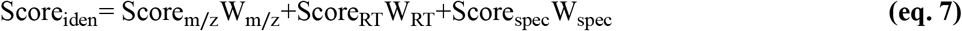

where Score_m/z_, W_m/z_ and W_RT_ are the same as in eq. 2. Score_RT_ is calculated as follows:

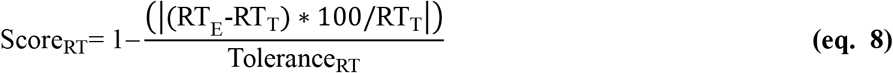

RT_E_ and RT_T_ are the experimental RT and theoretical RT, respectively. The Tolerance_RT_ represents the tolerance of RT match with a default value of 30%.

Score_spec_ is the MS2 spectral match score, which is scored using a dot-product function (eq. 1) with some modifications. If the *m/z* value of seed metabolite is larger than the neighbor metabolite, the fragment ions in the surrogate MS2 spectrum with *m/z* larger than that of the neighbor metabolite are removed. Vice versa, the fragment ions in the experimental MS2 spectrum with *m/z* larger than that of seed metabolite are also removed. W_spec_ is the weight of Score_spec_. The default values of W_m/z_, W_RT_ and W_spec_ are 0.25, 0.25 and 0.5, respectively. The identifications of each MS1 peak are ranked by Score_iden_. After the identification of neighbor metabolites, isotope peak annotation is also performed for each neighbor metabolite using the same procedures described above.

### Selection of seed metabolites and recursive identification

After one round of identification, new seed metabolites are selected from the identified neighbor metabolites to start recursive identification (Fig. 1c). Peaks with new identifications are selected as seed metabolites. MetDNA then repeats the neighbor metabolite identification and isotope peak annotation until there are no new seed metabolites available for the next round of annotation.

### Confidence assignment

MetDNA uses a multi-step strategy to evaluate the confidence of the metabolite identification. First, all the identified MS1 peaks are grouped according to their identification and RT. The MS1 peaks with the same metabolite identifications are grouped together and further divided into different peak groups according to their RT. The peak group is defined as a set of peaks (e.g., monoisotope peak, isotope peaks, and adduct peaks) with the same identification and in the same RT window (default is 3 seconds). If one peak has multiple identifications, it may belong to multiple peak groups. Similarly, one metabolite may also belong to multiple peak groups. The confidence is then assigned to each peak group and all MS1 peaks in the group according to the following rules:

1. Grade 1: at least one MS1 peak in the peak group is identified through the standard spectral library (or initial seed metabolite);
2. Grade 2: do not meet rule 1, and there are isotope peaks available in the peak group;
3. Grade 3: do not meet rules 1 and 2, and there are reliable adduct peaks in the peak group, such as [M+H]^+^, [M+Na]^+^ or [M+NH_4_]^+^ for positive mode, and [M−H]^−^, [M+Cl]^−^ or [M+CH_3_COO]^−^ for negative mode, respectively; and
4. Grade 4: the remaining peak groups that do not meet rules 1, 2, or 3.

### Redundancy removal

Identification redundancy includes peak redundancy and metabolite redundancy. Peak redundancy is defined as the total number of metabolite identifications divided by the total number of peaks with identifications, that is, the number of metabolites per peak. By contrast, the metabolite redundancy is defined as the total number of peak groups with identification divided by the total number of metabolite identifications, that is, the number of peak groups per metabolite. MetDNA then removes the identification redundancy according to the confidence levels of the peak group and the peaks in the group. First, if one metabolite matches multiple peak groups, the program removes the identification from all the peaks in the peak groups with grade 4. However, if all of the matched peak groups are grade 4, all the identifications are maintained. Second, if one peak matches multiple metabolites, the identifications with the highest grade are kept. After the removal of the identification redundancy, the constitution of the peak groups may change. The confidence assignment is then repeated, followed by a repetition of the redundancy removal process, which is also a recursive process. The recursive process continues until the identification redundancy remains unchanged. The identification redundancy is calculated as the mean value of the peak redundancy and the metabolite redundancy.

### Identification of dysregulated pathways

A pathway enrichment analysis is used to identify and characterize the dysregulated metabolic pathways. First, dysregulated peaks with identifications are selected according to a univariate test (Student’s *t*-test or Mann-Whitney-Wilcoxon test) with or without FDR correction. The maximum tolerance of *P*-values can be set by the users. A volcano plot is provided to demonstrate the distributions of the dysregulated peaks. Second, the metabolite identifications from the dysregulated peaks are mapped to the KEGG metabolic pathways. Currently, MetDNA contains the pathway information for 16 biological species (**Supplementary Data 10**). The hypergeometric test is used to evaluate whether the dysregulated metabolites are enriched in one pathway^19^. The *P*-value from the hypergeometric test is also calculated for each pathway. The dysregulated pathways with *P*-values less than 0.05 are output as dysregulated pathways. The information for dysregulated pathways is output into the file named “Pathway.enrichment.analysis”.

### Quantitative analysis of dysregulated pathway

MetDNA utilizes the quantitative information from the MS1 peaks to characterize the expression levels of the dysregulated pathways in a quantitative fashion. First, all the peak intensities are Pareto-scaled^24^. If one peak group has multiple MS1 peaks, then the most abundant peak is selected to represent the quantity of the peak group. Second, if one metabolite matches multiple peaks, then the peak with the highest identification score (Score_iden_) is selected to represent the metabolite. Third, the expression level of one dysregulated pathway is calculated as the average value of all the metabolites in the pathway. The quantitative results for the metabolites and pathways are output into two files named “Quantitative.pathway.metabolite.result” and “Quantitative.pathway.result”, respectively.

### Predicting the retention time

To obtain the theoretical retention times of all the metabolites in the MRN, MetDNA utilizes the quantitative structure-retention relationship (QSRR) to construct a prediction model to generate theoretical RTs^25, 26^. The RTs of metabolites under liquid chromatography (LC) highly depend on their structures and physiochemical properties, which can be described quantitatively using molecular descriptors (MDs)^25, 26^. With the QSRR prediction model, the input of a set of MD values for one metabolite could generate the theoretical RT. This approach requires the following data to establish a prediction model: (1) a training dataset containing a number of metabolites with experimental RTs; (2) a machine-learning-based algorithm; and (3) the MDs of metabolites. The detailed steps for the RT prediction are described below.

**Step 1. Calculating the molecular descriptors**. The R package “*rcdk*” is used to calculate the MDs of the metabolites from their SMILES structures. The SMILESs of metabolites from the in-house spectral library and the MRN were obtained using the Identifier Exchange Service of PubChem (https://pubchem.ncbi.nlm.nih.gov/idexchange/idexchange.cgi). A total of 346 MDs are calculated for each metabolite.
**Step 2. Obtain a training dataset**. The identified metabolites through the spectral match are used as the training dataset to establish the prediction model. If one MS1 peak has multiple metabolite identifications, or vice versa, then the unique metabolite identification is selected using the following criteria: (1) if one MS1 peak has multiple identifications, the one with the highest DP score is kept; (2) if one metabolite matched to multiple MS1 peaks, then the one with the highest intensity is kept; and (3) metabolite identifications with reliable adducts, such as [M+H]^+^, [M+Na]^+^ and [M+NH_4_]^+^ in positive ionization and [M−H]^−^, [M+CH3COO]^−^ and [M+Cl]^−^ in negative ionization are selected. For the metabolites in the training dataset, MDs with more than 50% missing values (MVs) across all metabolites are removed. The remaining MVs in the training dataset are imputed using the KNN algorithm (with the R package “*impute*”). The MDs with the same values across all the metabolites are also removed. As a result, a training dataset is obtained, including a set of metabolites with experimental RTs, and the MDs for the metabolites are calculated.
**Step 3. Optimize the combination of MDs for prediction**. A machine-learning-based algorithm known as the random forest (RF) is used to develop the prediction model. First, the combination of MDs is optimized for prediction. An RF model is constructed using MDs as independent variables and RTs as dependent variables (with the R package “*randomForesť*). Approximately 68% of the metabolites are randomly selected as the “internal training dataset”, and the remaining metabolites are used as the “internal test dataset”. An “importance” value is assigned to each MD to evaluate its contribution to the prediction model. The model construction is repeated 100 times. Each time, the top 5 ranked MDs according to the importance values are recorded. The MDs that which appear > 50 times in 100 RF models are selected as the optimized MD combination. Using the datasets in our lab, we optimized two combinations of MDs for HILIC and reverse phase (RP) separations, respectively. The optimized combination of MDs for the HILIC includes XLogP, tpsaEfficiency, WTPT.5, khs.dsCH, MLogP, nAcid, nBase and BCUTp.1l. The optimized combination of MDs for the RP phase includes XLogP, WTPT.5, WTPT4, ALogp2 and BCUTp.1l.
**Step 4. Parameter optimization**. The parameters in the RF algorithm, ntree (i.e., number of trees to grow) and mtry (i.e., number of variables randomly sampled as candidates at each split) are also optimized. The two parameters are combined together to form a set of parameter combinations. The performance of each parameter combination is evaluated using the mean squared error (MSE). The parameter combination with the smallest MSE is used to construct the final prediction model.
**Step 5. Retention time prediction**. With the RF-based prediction model, the theoretical RTs of all the metabolites in the spectral library and MRN are obtained using their calculated MDs. The theoretical RTs are used to improve the confidence in the metabolite identifications.

### Standard MS2 spectral library

The standard MS2 spectral library is used to identify the initial seed metabolites in MetDNA. The curation of the library is provided in our previous publication^27^. All the MS2 spectra were acquired on Sciex TripleTOF 5600 or 6600 instruments with commercial metabolite standards. For each metabolite, the targeted product ion scans were applied to acquire the MS2 spectrum with a flow injection method. The curation of the spectral library follows the instructions and protocols in a publication from the NIST to improve spectral reproducibility^28^. In brief, for each metabolite, at least 11 MS2 spectra were acquired. The cluster of MS2 spectra with high similarities (DP > 0.7) was selected to generate a consensus MS2 spectrum. MS2 spectra at different levels of collision energy (10, 20, 30, 40, and 35 ± 15) were acquired. The current library in MetDNA contains 841 metabolites in total, with 841 for the positive mode and 837 for the negative mode (**Supplementary Data 11**).

### Validation of metabolite identification

A validation strategy was designed to validate the metabolite identifications from MetDNA (**Supplementary Fig. 11**). First, the chemical structures of initial seed metabolites are confirmed using metabolite standards with a combination of *m/z* match (*m/z* error < 25 ppm), MS2 spectral match (DP > 0.8) and RT match (RT error < 60 s). These chemical structures are considered as Level 1 identifications according to MSI, and referred as “true structures”. Then, 30% of initial seed metabolites are randomly selected and used as the seed metabolites for MRN-based recursive metabolite identification. The remaining 70% of initial seed metabolites having true structures are used as the set of validation metabolites. The MetDNA identification results from the 30% of initial seed metabolites are compared to the set of validation metabolites. For each MS1 peak in the validation set, the identifications from MetDNA are classified as “correct”, “isomer” and “error”. Specifically, if one of top 3 identifications from the MetDNA is the true structure, the identification is denoted as “correct”. If one of top 3 identifications from MetDNA is the isomer of true structure, the identification is denoted as “isomer”. If none of top 3 identifications from MetDNA is isomer of true structure, the identification is denoted as “error”. In MetDNA, the identifications are ranked by their Score_iden_. The random selection of 30% of initial seeds repeats 10 times, and the results are summarized in **Supplementary Data 1** for aging fruit fly (dataset #1) and **Supplementary Data 2** for *E. coli* (dataset #9).

### Reagents, fruit fly culture and sample preparation

LC-MS grade water (H_2_O) and methanol (MeOH) were purchased from Honeywell (Muskegon, USA). LC-MS grade acetonitrile (ACN) was purchased from Merck (Darmstadt, Germany). Ethanol was purchased from Sinopharm (Beijing, China). Ammonium hydroxide (NH_4_OH) and ammonium acetate (NH_4_OAc) were purchased from Sigma-Aldrich (St. Louis, USA). Metabolite chemical standards were purchased from J&K (Beijing, China), Sigma (St. Louis, USA), Carbosynth (Berkshire, UK), TCI (Tokyo, Japan) and Energy Chemical (Shanghai, China).

Wild-type male fruit flies (FlyBase ID: FBst0005905) were cultured in standard media (temperature, 25 °C; humidity, 60%; and a 12 h light and 12 h dark cycle). At day 3 (3-day) and day 30 (30-day), 100 fruit flies were collected and divided into 10 samples (10 flies in each sample, n = 10 in each group, and 20 samples in total). The fruit flies were killed with 75% ethanol. The heads of the fruit flies were collected and placed into microcentrifuge tubes. The tubes were immediately frozen with liquid nitrogen and stored at −80 °C until metabolite extraction. The fruit fly samples were defrosted on ice. The fruit fly samples were then homogenized with 200 μL of H_2_O and 20 ceramic beads (diameter, 0.1 mm) using a homogenizer (Precellys 24, Bertin Technologies). A mixture of ACN:MeOH (1:1, v/v; 800 μL) was added to the samples, which were then vortexed for 30 s, followed by incubation in liquid nitrogen for 1 min, and then thawed on ice. This vortex-freeze-thaw cycle was repeated three times. The samples were incubated for 1 hour at −20 °C for protein precipitation, followed by centrifugation at 13,000 rpm and 4 °C for 15 min. The supernatant solution was removed and evaporated to dryness in a vacuum concentrator (Labconco, USA). A mixture of ACN:H_2_O (1:1, v/v; 100 μL) was then added to reconstitute the dry extracts, followed by sonication (50 Hz, 4 °C) for 10 min. The solutions were centrifuged at 13,000 rpm and 4 °C for 5 min to precipitate the insoluble debris. Finally, the supernatant solutions were transferred to HPLC glass vials and stored at −80 °C prior to LC-MS/MS analysis.

### LC-MS/MS analysis of fruit fly samples

The metabolomics data acquisition for fruit fly samples was performed using a UHPLC system (1290 series, Agilent Technologies, USA) coupled to a quadruple time-of-flight mass spectrometer (TripleTOF 6600, AB SCIEX, USA). A Waters ACQUITY UPLC BEH Amide column (particle size, 1.7 μm; 100 mm (length) × 2.1 mm (i.d.)) was used for the LC separation and the column temperature was kept at 25 °C. Mobile phase A was 25 mM ammonium hydroxide (NH_4_OH) + 25 mM ammonium acetate (NH_4_OAc) in water, and B was ACN for both the positive (ESI+) and negative (ESI−) modes. The flow rate was 0.3 mL/min and the gradient was set as follows: 0-1 min: 95% B, 1-14 min: 95% B to 65% B, 14-16 min: 65% B to 40% B, 16-18 min: 40% B, 18-18.1 min: 40% B to 95% B and 18.1-23 min: 95% B. The injection volume was 2 μL. All the samples were randomly injected during data acquisition.

The data acquisition was operated using the information-dependent acquisition (IDA) mode. The source parameters were set as follows: ion source gas 1 (GAS1), 60 psi; ion source gas 2 (GAS2), 60 psi; curtain gas (CUR), 30 psi; temperature (TEM), 600 °C; declustering potential (DP), 60 V or −60 V in positive or negative modes, respectively; and ion spray Voltage floating (ISVF), 5500 V or −4000 V in positive or negative modes, respectively. The TOF MS scan parameters were set as follows: mass range, 60-1200 Da; accumulation time, 200 ms; and dynamic background subtract, on. The product ion scan parameters were set as follows: mass range, 25-1200 Da; accumulation time, 50 ms; collision energy (CE), 30 V or −30 V in positive or negative modes, respectively; collision energy spread (CES), 0; resolution, UNIT; charge state, 1 to 1; intensity, 100 cps; exclude isotopes within 4 Da; mass tolerance, 10 ppm; maximum number of candidate ions to monitor per cycle, 6; and exclude former target ions, for 4 seconds after 2 occurrences.

### Data processing of the fruit fly dataset

All 20 MS raw data files (.wiff) were separately converted to mzXML format and mgf format using ProteoWizard (version 3.0.6150, http://proteowizard.sourceforge.net/). The detailed parameters for the data conversion are listed in Table 1. First, the mzXML data files were grouped into two folders (named “W03” and “W30”) and subjected to peak detection and alignment using the R package called “*xcms*” (version 1.46.0, https://bioconductor.org/packages/3.2/bioc/html/xcms.html)^17^. The detailed code for XCMS processing is provided in **Supplementary Note 2**. The key parameters were set as follows: method = “centWave”; ppm = 15; snthr = 10; peakwidth = c(5, 40); minifrac = 0.5. The generated MS1 peak table includes the mass-to-charge ratio (*m/z*), retention time (RT), peak abundances, and other information. The peak table was then modified as follows: (1) for the first 12 columns, those named “name”, “mzmed” and “rtmed” were kept, and the others were deleted; (2) the first three columns were renamed “name”, “mz” and “rt”. The generated MS1 peak tables (one for positive mode and one for negative mode) are used for the MetDNA analysis.

**Table 1.** The detailed parameters for data conversion using ProteoWizard.

| Parameters | mzXML conversion | mgf conversion |
| --- | --- | --- |
| **Output format** | mzXML | mgf |
| **Binary encoding precis** | 64-bit | 64-bit |
| **Write index** | YES | YES |
| **Use zlib compression** | YES | YES |
| **TPP compatibility** | YES | YES |
| **Package in gzip** | NO | NO |
| **Use numpress linear compression** | NO | NO |
| **Use numpress short logged floate compression** | NO | NO |
| **Use numpress short positive inter compression** | NO | NO |
| **Filters** | Peak Picking | Peak Picking |

Second, a “sample information” file (.csv format) is prepared to describe the sample group information. The first column is named “sample.name”, while the second one is named “group”. Two group names (“W03” and ‘W30”) are provided.

Finally, the MS1 peak table, sample information file, and MS2 data files (.mgf format) were all uploaded to our MetDNA webserver (http://metdna.zhulab.cn) for data analysis. Positive and negative datasets were processed together. The data processing parameters for MetDNA were set as follows: Ionization polarity, Both; Liquid Chromatography, HILIC; MS Instrument, Sciex TripleTOF; Collision Energy, 30; Control Group, W03; Case Group, W30; Univariate Statistics, Student’s *t*-test; Species, *Drosophila melanogaster* (fruit fly); Cutoff of *P*-value, 0.01; and *P*-value Adjustment, Yes. The generated analysis result is provided in **Supplementary Data 12**. The detailed code for XCMS processing, parameter settings for MetDNA are provided in **Supplementary Data 13**.

### Transcriptomics data for the aging fruit fly

The RNA-seq of the aging fruit fly head tissues (3-day *vs*. 30-day, n = 3 for each group) was obtained from our recent study^29^ and downloaded from the Gene Expression Omnibus (https://www.ncbi.nlm.nih.gov/geo/, GEO: GSE96654). In brief, the head tissue samples of the fruit flies were sequenced using Illumina NextSeq 550 or Hisep 2500 platforms with single end 100 bps. The sequencing reads were mapped to the reference genome dm6 with STAR2.3.0. The read counts for each gene were calculated using HTSep-0.5.4. The count files were normalized with the R package “*DESeq*”. Finally, the FlyDatabase ID for each gene was transformed to KEGG ID with the R package “*clusterProfiler*”. The genes without mapped KEGG IDs were removed from the dataset. The final transcriptomics dataset of aging fruit flies is provided in **Supplementary Data 7**.

### Multi-omics integration of metabolomics and transcriptomics

A correlation network between the metabolomics and transcriptomics was constructed^20^. To construct the correlation network at the pathway level, we first obtained quantitative pathway data from the metabolomics data (see **Quantitative analysis of dysregulated pathway**). For the transcriptomics data, the same method was applied. In brief, genes in the same pathway were grouped, and the mean value of the genes was then taken for each pathway to represent the quantitative information of this pathway. For pathways in the metabolomics and transcriptomics data, all the pairwise Pearson correlations were calculated to construct the correlation network at the pathway level for both types of data (Student’s-*t* test, *P*-values < 0.05, only the same pathways in both types of data). The correlation network at the gene and metabolite level for each pathway was also constructed using the same method with a Pearson correlation (Student’s-*t* test, *P*-values < 0.01 and absolute Pearson correlation values > 0.7). The network was visualized using Cytoscape (version 3.2.1, http://www.cytoscape.org/).

### Software availability

MetDNA was developed using a mixture of R, JavaScript, and Python and is available for non-commercial use at http://metdna.zhulab.cn/. The webserver is currently hosted on a Linux server from Alibaba Cloud (https://www.alibabacloud.com/) with 8 cores (3.2 GHz CPUs) and 16 GB RAM. A help document for using MetDNA can be found at http://metdna.zhulab.cn/help. The demo data is provided to learn how to use MetDNA at http://metdna.zhulab.cn/demo.

### Data availability

The metabolomics datasets of aging fruit fly can be accessed at MetaboLights (Project ID: MTBLS612 for positive and MTBLS615 for negative modes, respectively). The metabolomics datasets of mouse liver tissues can be accessed at MetaboLights (Project ID: MTBLS601 for positive and MTBLS606 for negative modes, respectively).

---

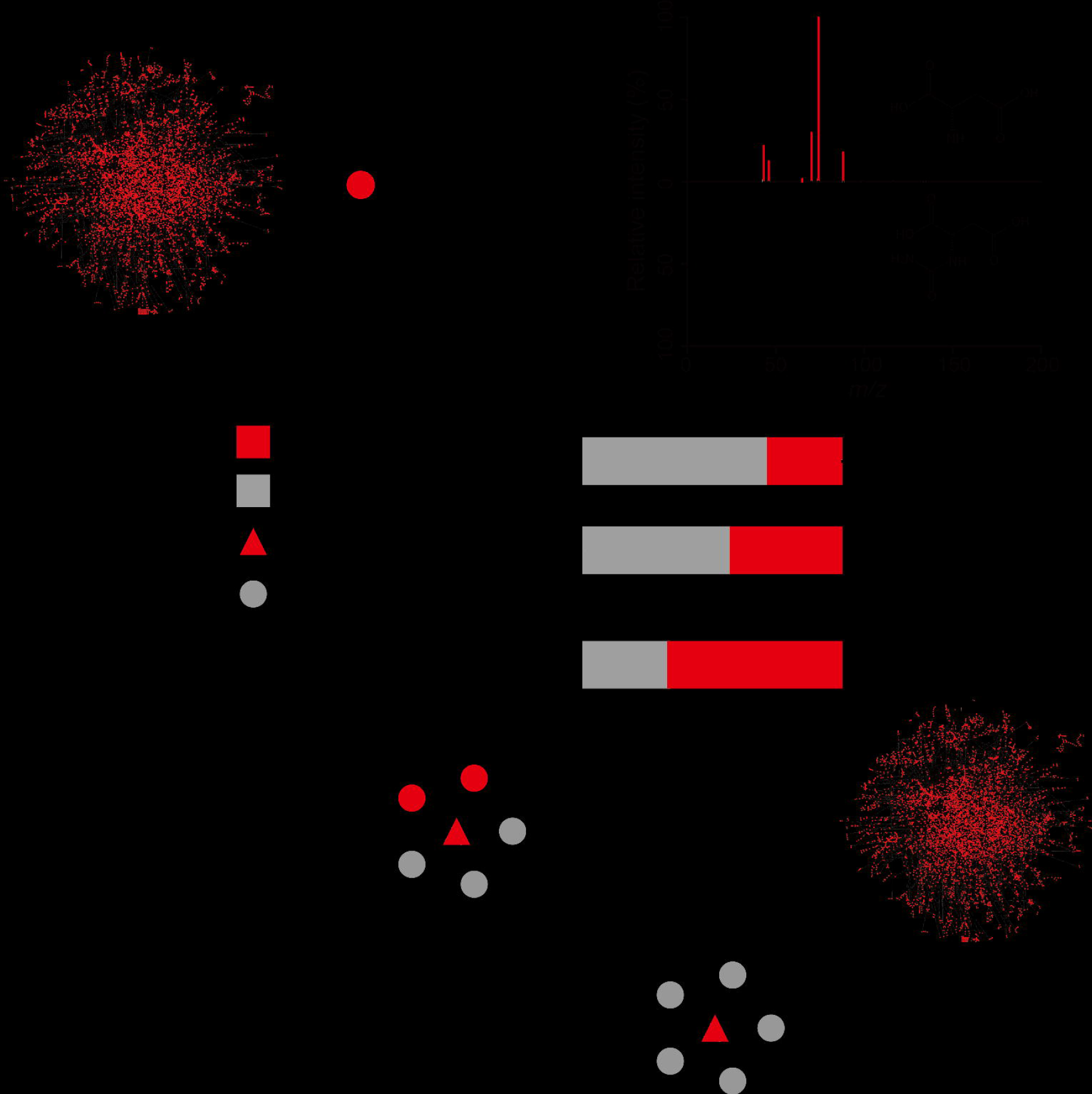

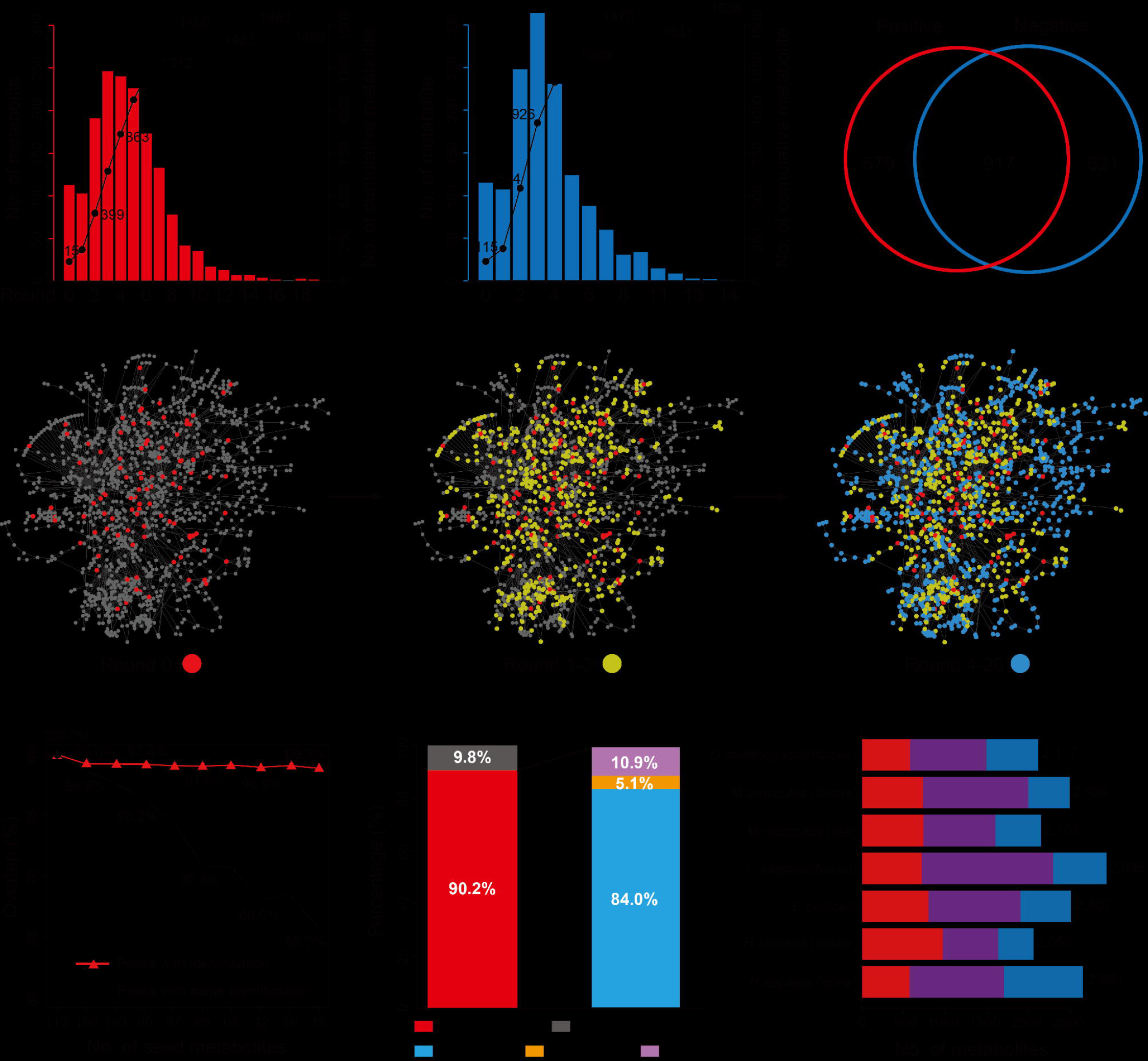

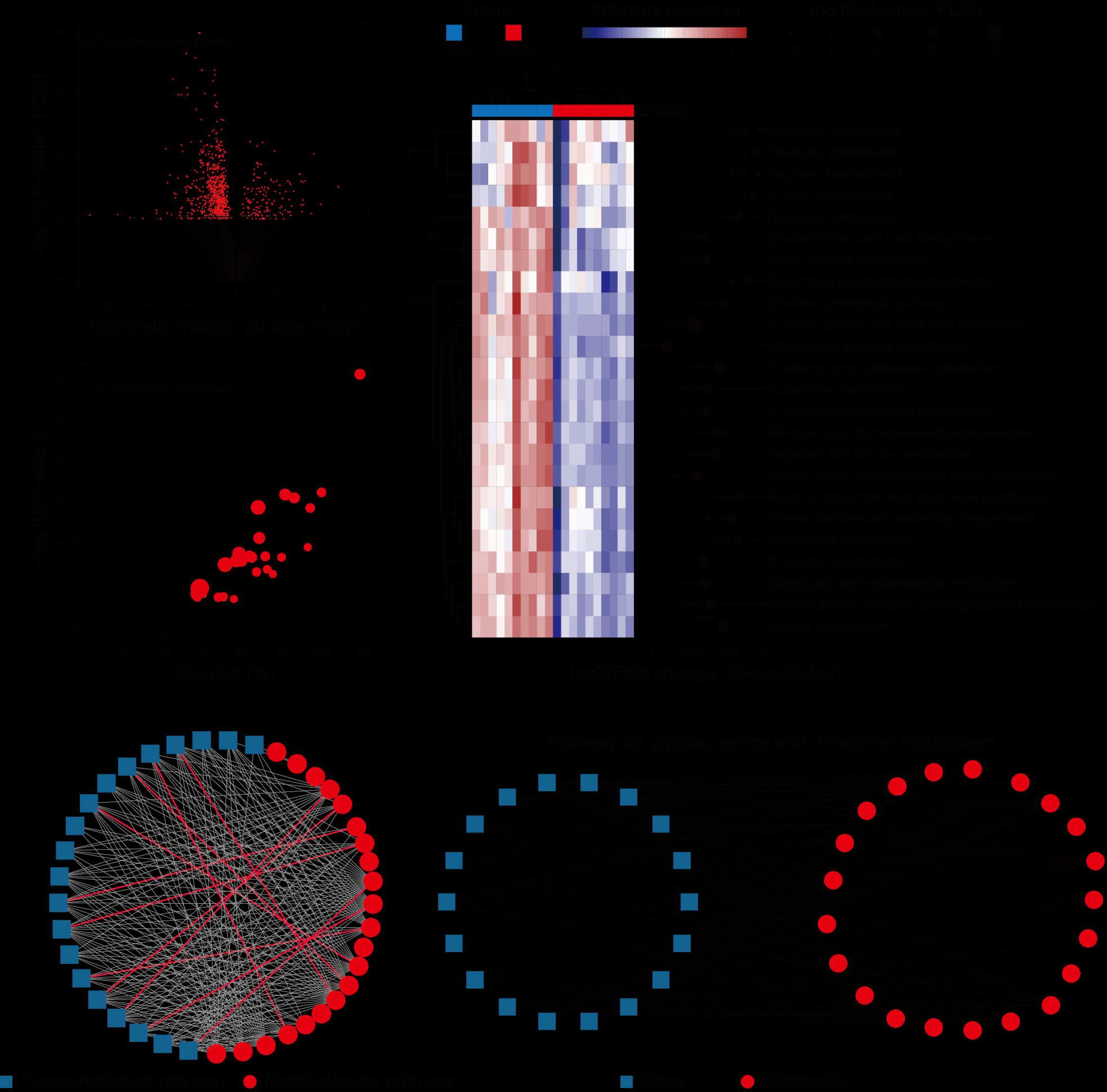

## REFERENCES

1. Fiehn O. Plant Mol. Biol. 48, 155–171 (2002).

2. Nicholson J.K. & Lindon J.C. Nature 455, 1054–1056 (2008).

3. Patti, G.J., Yanes, O. & Siuzdak, G. Nat. Rev. Mol. Cell Biol. 13, 263–269 (2012).

4. Johnson, C.H., Ivanisevic, J. & Siuzdak, G. Nat. Rev. Mol. Cell Biol. 17, 451–459 (2016).

5. Wishart D.S. Nat. Rev. Drug Discov. 15, 473–484 (2016).

6. Zhu Z.-J. et al. Nat. Protoc. 8, 451–460 (2013).

7. Wishart D.S. et al. Nucleic Acids Res. 41, D801–807 (2013).

8. Smith C.A. et al. Ther. Drug Monit. 27, 747–751 (2005).

9. Vinaixa M. et al. TrAC Trends Anal. Chem. 78, 23–35 (2016).

10. Wolf S. et al. BMC bioinformatics 11, 148 (2010).

11. Duhrkop K. et al. P. Natl. Acad. Sci. 112, 12580–12585 (2015).

12. Huan T. et al. Anal. Chem. 87, 10619–10626 (2015).

13. Allen, F., Greiner, R. & Wishart, D. Metabolomics 11, 98–110 (2014).

14. Tsugawa H. et al. Anal. Chem. 88, 7946–7958 (2016).

15. Blazenovic I. et al. J. Cheminformatics 9, 32 (2017).

16. Kanehisa M. et al. Nucleic Acids Res. 42, D199–205 (2014).

17. Smith C.A. et al. Anal. Chem. 78, 779–787 (2006).

18. Sumner L.W. et al. Metabolomics 3, 211–221 (2007).

19. Xia J. & Wishart D.S. Bioinformatics 26, 2342–2344 (2010).

20. Li S. et al. Cell 169, 862–877 (2017).

## REFERENCES

21. Tsugawa H. et al. Nat. Methods 12, 523–526 (2015).

22. Stein S.E. & Scott D.R. J. Am. Soc. Mass Spectr. 5, 859–866 (1994).

23. Kuhl C. et al. Anal Chem. 84, 283–289 (2012).

24. van den Berg, R.A. BMC genomics 7, 142 (2006).

25. Wolfer A.M. et al. Metabolomics 12 (2015).

26. Tsugawa H. et al. J. of Cheminformatics 9, 19 (2017).

27. Li H. et al. Anal. Chem. 88, 8757–8764 (2016).

28. Yang, X., Neta, P. & Stein, S.E. Anal. Chem. 86, 6393–6400 (2014).

29. Ma Z. et al. bioRxiv doi: https://doi.org/10.1101/247726 (2018).

